# Analytical solution to the Flory-Huggins model

**DOI:** 10.1101/2022.04.23.489256

**Authors:** Daoyuan Qian, Thomas C. T. Michaels, Tuomas P. J. Knowles

## Abstract

A self-consistent analytical solution for binodal concentrations of the two-component Flory-Huggins model is derived. We show that this form extends the validity of the Ginzburg-Landau expansion away from the critical point to cover the whole phase space. Furthermore, this analytical solution reveals an exponential scaling law of the dilute phase binodal concentration as a function of interaction strength and chain length. We demonstrate explicitly the power of this approach by fitting experimental protein liquid-liquid phase separation boundaries to determine the effective chain length and the solute-solvent interaction energy. Moreover, we demonstrate that this strategy allows us to resolve the differences in the interaction energy contributions of individual amino acids. This analytical framework can serve as a new way to decode the protein sequence grammar for liquid-liquid phase separation.

## I. INTRODUCTION

The formation of membraneless organelles through liquid-liquid phase separation (LLPS) has emerged as an important mechanism used by cells to regulate their internal biochemical environments, and it is also closely related to development of neurodegenerative diseases [1–8]. The Flory-Huggins model [9–11] is a foundational theoretical picture that describes the phenomenology of LLPS, driven by a competition of entropy and interaction energy. Despite the generality of the Flory-Huggins model, analytical solutions describing the binodal line have not been available, and progress has instead been made through numerical methods [11–13]. Here we propose an analytical self-consistent form for the binodal concentrations and demonstrate the high accuracy comparable to numerical schemes. This can then be used to efficiently fit experimental binodal data and determine key underlying physical parameters.

## II. THE FLORY-HUGGINS MODEL FOR BINARY MIXTURES

The two-component Flory-Huggins theory describes mixing of a polymer species of length *N* with a homogeneous solvent. Denoting the volume fraction of polymers as *ϕ*, the volume fraction of the solvent is simply 1 *− ϕ* by volume conservation. The model uses an effective lattice site contact energy 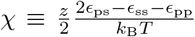 between polymer and solvent in which *z* is a coordination constant, and *ϵ*_ps_, *ϵ*_ss_ and *ϵ*_pp_ are bare polymer-solvent, solvent-solvent and polymer-polymer contact energies. The free energy density of the Flory-Huggins model is given by [9–12]

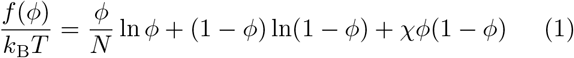

where *k*_B_ is the Boltzmann constant and *T* the absolute temperature. In the following we consider energies relative to the thermal energy and set *k*_B_*T* = 1 to simplify notation. The first two terms on the right hand side of equation (1) represent the entropic energy of mixing while the third term denotes the effective contact energy. Two important quantities can be calculated: the spinodal concentration and binodal concentration. The spinodal is the boundary between locally stable/unstable regions and can be solved exactly, while the binodal separates globally stable/unstable regions and the system can still be locally stable on the binodal boundary itself. It is also straightforward to generalise equation (1) to include more components or surface tension [11, 14, 15], and over the decades more detailed models have been proposed to include electrostatic interactions [16], sticker-spacer behaviours [7, 8, 17] or to calculate free energy density from first principle using a field-theoretic approach [18–21]. It thus appears that the Flory-Huggins theory is an outdated model due to its over-simplifying, mean-field nature, while we note that even then an analytical solution for the binodal concentrations is lacking for this most basic picture of LLPS.

### Spinodal concentrations

The free energy density becomes locally unstable at *f*″(*ϕ*) ≤ 0, and consequently the spinodal boundary *ϕ*^spi^ is defined at the transition point *f*″(*ϕ*^spi^) = 0. Solving for this condition we obtain the dense 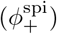 and dilute phase 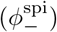 spinodal concentrations

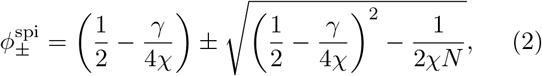

with 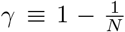, which goes to zero in the symmetric *N* = 1 case. The critical point of LLPS occurs when the dense and dilute phases coincide, corresponding to a critical interaction strength *χ*_c_ and concentration *ϕ*_c_:

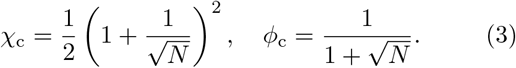

Near the critical point with *δχ* ≡ *χ − χ*_c_ ≈ 0 the spinodal concentrations have the approximate form

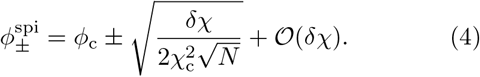

Note that in the opposite limit of large *N* or large *χ* the dilute phase concentration has a power-law scaling

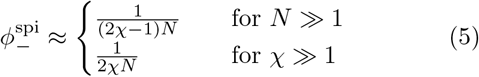

and these are qualitatively different from the exponential scaling of the dilute phase binodal concentrations, as will be shown using the self-consistent equations.

### Binodal concentrations

The binodal concentrations are found by assuming the existence of two distinct phases characterised by polymer volume fractions *ϕ*_+_, *ϕ*_−_, and phase volumes *V*_+_, *V*_−_. Equilibrium condition requires minimisation of the total energy *F*_tot_ ≡ *V*_+_*f* (*ϕ*_+_) + *V*_−_*f* (*ϕ*_−_) subject to total volume and mass conservation conditions *V*_+_ + *V*_−_ = *V*_tot_ and *V*_+_*ϕ*_+_ + *V*_−_*ϕ*_−_ = *V*_tot_*ϕ*_tot_. Using Lagrange minimisation we identify the chemical potential *μ*(*ϕ*) ≡ *f*′(*ϕ*) and osmotic pressure Π(*ϕ*) ≡ *ϕf*′(*ϕ*) − *f* (*ϕ*) as Lagrange multipliers that have to hold the same values in the two compartments:

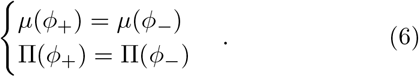

Graphically, in a [*ϕ, f* (*ϕ*)] plot, the *μ*(*ϕ*_+_) = *μ*(*ϕ*_−_) condition forces the two points describing the co-existing phases to have the same gradient and Π(*ϕ*_+_) = Π(*ϕ*_−_) aligns the two tangent lines to have the same y-intercept, and as such represent a common tangent construction (figure 1A). Using the Flory-Huggins expression (1) we have

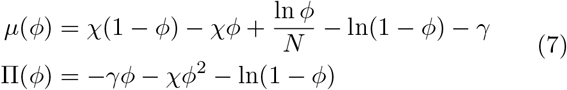

and the binodal concentrations can be calculated by solving (6) with the definitions of (7). The objective of this paper is to generate analytical solutions of equation (7). As a first step, an approximate binodal solution near the critical point can be worked out by expanding the free energy around *ϕ* = *ϕ*_c_ + *δϕ* and *χ* = *χ*_c_ + *δχ* with small *δχ* and *δϕ*. Terms that are constant or linear in *δϕ* drop out of the common tangent construction; terms of order higher than 4 are also truncated. The result is 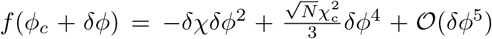 and for *δχ >* 0 this is a simple Ginzburg-Landau second order phase transition. The binodal concentrations near the critical point are

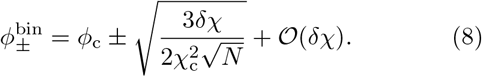

The Ginzburg-landau solution describes the binodal near the critical point (figure 1B, C). We now aim to extend this solution to cover *χ* far away from *χ*_c_ through a self-consistent approach, summarised as the following. Suppose we need to solve the equation *η* = *𝒜*(*η*) with some operator *𝒜*. Instead of solving it directly, we treat *𝒜* as a discrete map and start with a solution *η*^(0)^ and apply *𝒜* iteratively to generate the orbit *η*^(*i*)^ ≡ *𝒜*^*i*^[*η*^(0)^] with *i* = 1, 2, …. With a suitable form of *𝒜* the orbit converges to the fixed point lim_*i*→∞_ *η*^(*i*)^ = *η*, which then solves the equation *η* = *𝒜* (*η*). This is the contractive mapping principle [22, 23]. The self-consistent approach has been previously employed to approximate the protein aggregation kinetics curves [24, 25] and here we show a similar procedure allows efficient and accurate computation of the binodal concentrations.

**FIG. 1.**
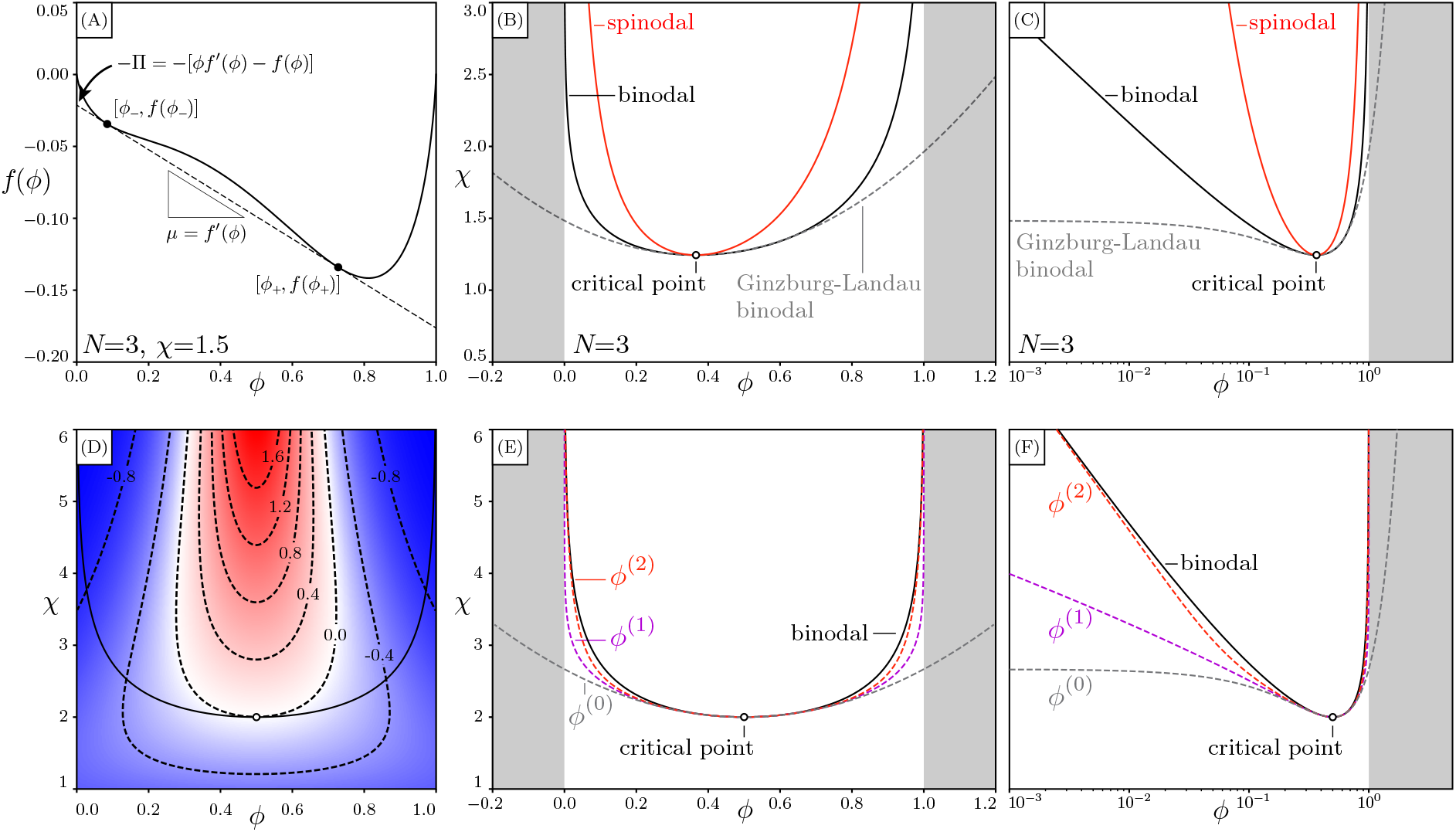
Flory-Huggins model (A) - (C) and self-consistent solution for the symmetric *N* = 1 case (D) - (H). (A) Common tangent construction at *N* = 3, *χ* = 1.5 gives the dense and dilute phase concentrations *ϕ*_1,2_. The gradient of the common tangent is the chemical potential *μ*(*ϕ*) and the y-intercept is 1 times the osmotic pressure Π(*ϕ*). (B), (C) Complete phase diagram of *N* = 3 system in linear and logarithmic *ϕ* scales. The binodal is calculated numerically [11]. Near the critical point the Ginzburg-Landau binodal approximates the exact binodal well, but at large *χ* the two quickly diverges and the Ginzburg-Landau solution enters the unphysical range *ϕ <* 0 and *ϕ >* 1 (grey zones). (D) Plot of |*g*′(*ϕ*)| − 1 in *ϕ, χ* space. Black solid line is binodal and hollow circle is the critical point. Dashed lines are contours of constant |*g*′(*ϕ*)| − 1. The blue region with |*g*′(*ϕ*)| − 1 < 0 has stable orbits while red regions are unstable. (E), (F) Comparison between the numerical binodal (black solid line) with self-consistent schemes with 0, 1 and 2 iterations (coloured dashed lines).

## III. SELF-CONSISTENT SOLUTION FOR *N* = 1

### Self-consistent equation

With unit polymer length *N* = 1 the free energy (1) is symmetrical, and the binodal is given by the condition *f*′(*ϕ*) = 0, leading to 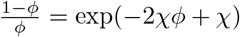. Upon rearrangement the binodal equation is

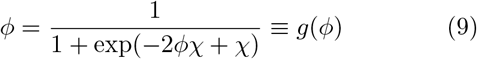

and we use *g*(*ϕ*) to define the 1D map

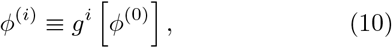

with the initial guess *ϕ*^(0)^ to be determined later. The fixed points are the binodal concentrations 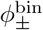. To study the convergent properties of the map we expand *g*(*ϕ*) near the fixed point, writing 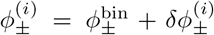 [26]. This gives 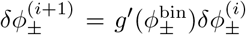 so for a convergent orbit we require 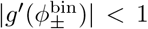, and quick convergence can be expected for *g* 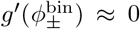. We calculate |*g*′(*ϕ*) | − 1 in the *ϕ, χ* space and observe that near the binodal, convergence is fast in the high-*χ* regime with |*g*′(*ϕ*) | ≈ 0 while much slower near criticality *χ* ≈ *χ*_c_ = 2, and becomes 0 at exactly the critical point (figure 1D). We thus need the initial guess to be close to the binodal just at *χ* ≈ *χ*_c_ and both the spinodal and approximate binodal may seem to be appropriate choices. It is worth writing down these concentrations near criticality with 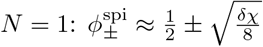 and 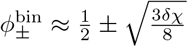. Furthermore, we can also obtain the approximate form of the |*g*′(*ϕ*)| − 1 = 0 contour near *χ*_c_ = 2 and 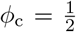, which gives exactly 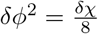. The spinodal thus coincides with the metastable line and is not an appropriate choice, despite it having better behaviour than the approximate binodal at large *χ*: the latter enters the unphysical regions *ϕ <* 0 and *ϕ >* 1 while the spinodal is always bound in 0 *< ϕ <* 1. We thus use the Ginzburg-Landau bimodal (8) as the initial guess so

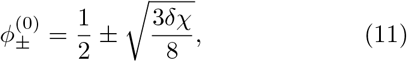

and the contraction mapping principle allows us to write the solution as

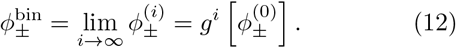

Good convergence is observed within two iterations (figure 1E, F).

### First order analytical solution

Performing one self-consistent step we obtain

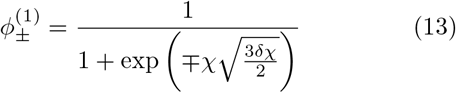

and this already improves the Ginzburg-Landau solution in linear *ϕ* scale. However, the large-*χ* behaviour is poorly captured in log scale due to the initial guess entering the unphysical region *ϕ <* 0 and *ϕ >* 1, and this problem disappears when a second self-consistent step is performed.

### Second order analytical solution

Performing one more self-consistent step we arrive at

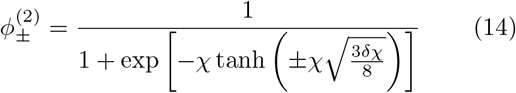

These expressions cover both the low and high-*χ* regime (figure 1 E, F). For large *χ ≫* 1 we have tanh 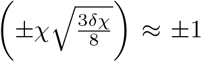, thus giving the scaling law for dilute phase binodal concentration

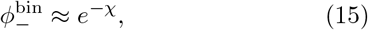

which is qualitatively different from the polynomial scaling of the spinodal concentration 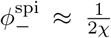. This exponential scaling has a physical interpretation as the chemical potential in the dilute limit takes the form of *μ* ≈ ln *ϕ*. An important implication is thus that the binodal phase separation can occur over a concentration range spanning orders of magnitude, while spinodal decomposition has a much narrower band of concentrations.

## IV. SELF-CONSISTENT SOLUTION FOR GENERAL *N*

### Self-consistent equation

Here we extend the *N* = 1 solution to the general case. Using (6) with (7) the binodal equations are:

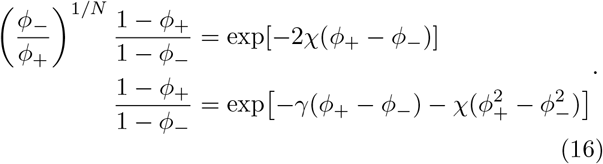

Define the two exponents

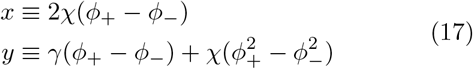

equations (16) can be simplified to *ϕ*_−_*/ϕ*_+_ = *e*^*−N*(*x−y*)^. Substituting this back into the second equation of (16) we get 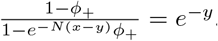. Solving for *ϕ*_+_ and then *ϕ*_−_ we obtain

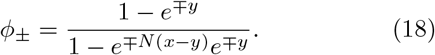

Organising *ϕ*_+_, *ϕ*_−_ in vector form:

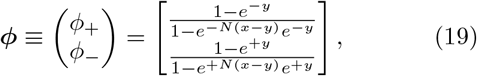

define the operator on the right hand side of (19) as ***G*** we have the map

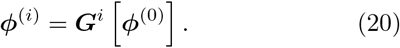

with the Ginzburg-Landau solution as the initial guess

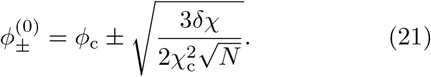

A scaling law for the dilute phase at large interaction strength can be obtained as before. At large *χ* we approximately have 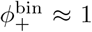 and 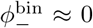, giving 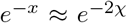 and *e*^*−y*^ ≈ *e*^*−γ−χ*^. The self-consistent expression for *ϕ*_−_ then becomes 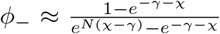. This can be further approximated to be

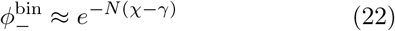

In the case of *N* = 1 we recover the *e*^*−χ*^ scaling discussed above. The convergence behaviour of the 2D map can also be studied similar to the 1D case. Define the Jacobian matrix ***J*** as

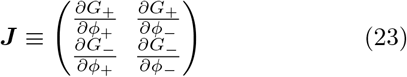

and writing ***ϕ***^(*i*)^ = ***ϕ***^bin^ + *δ****ϕ***^(*i*)^ we get *δ****ϕ***^(*i*+1)^ = ***J*** *δ****ϕ***^(*i*)^. Stability requires moduli of eigenvalues of ***J*** to be less than 1. Since both the eigenvalues and eigenvectors of ***J*** are in general complex, to better visualise the convergence of the orbit ***ϕ***^(*i*)^ we instead promote the discrete map to a continuous flow equation parametrised by *t*: ***ϕ***(*t*), with 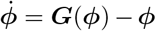. The velocity field 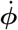 then contains the behaviour of the orbit ***ϕ***^(*i*)^ in the limit of small time steps and 3 fixed points can be identified: 1 stable fixed point corresponding to the binodal and 2 unstable fixed points on the *ϕ*_+_ = *ϕ*_−_ diagonal (figure 2A). We observe the orbit ***ϕ***^(*i*)^ is indeed convergent (figure 2B, C).

**FIG. 2.**
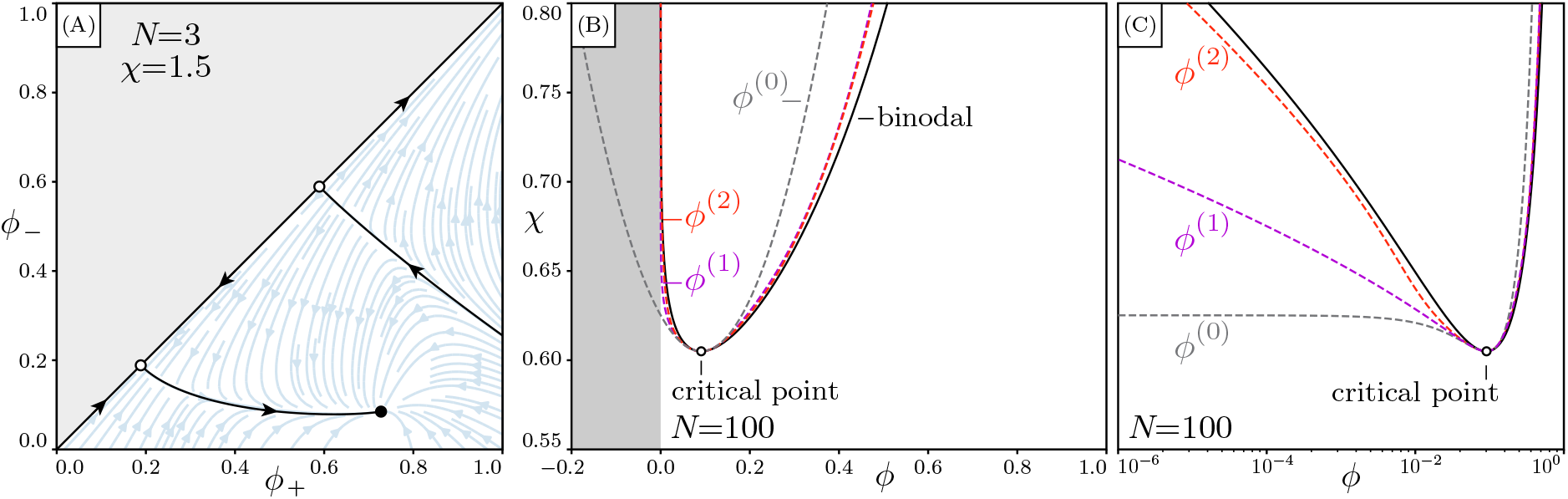
Self-consistent solution for general *N*. (A) Flow field of the continuous map with *N* = 3, *χ* = 1.5. Solid circle is the stable fixed point corresponding to the binodal, and hollow circles are saddle points. (B), (C) Comparison between the numerical binodal (black solid line) with self-consistent schemes with 0, 1 and 2 iterations (coloured dashed lines).

### First order analytical solution

To simplify notations we define two parameters

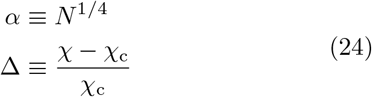

and express all other parameters in terms of *α*, Δ. This allows us to write

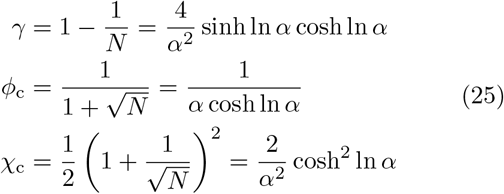

The Ginzburg-Landau solution [equation (21)] then takes the simple form

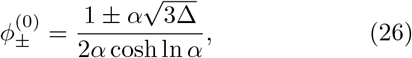

and *x, y* as defined in equation (17) are now

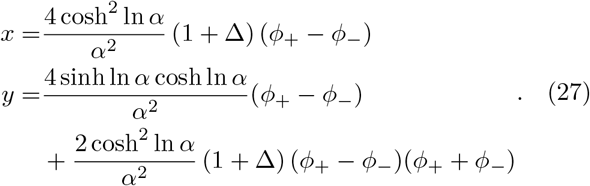

Direct substitution gives

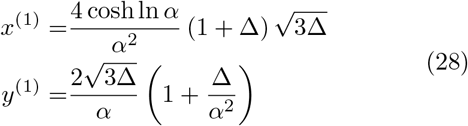

so at one self-consistent step we have

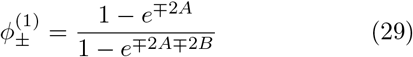

where

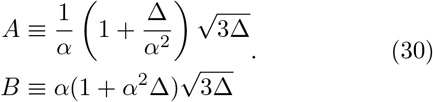

*A* and *B* are related through the transformation 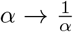 as 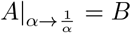 and vice versa. The large-*χ* behaviour is incompletely captured in log scale (figure 2C). We thus again calculate the second-order self-consistent solution.

### Second order analytical expression

At second order we substitute equation (29) into equation (27) and obtain

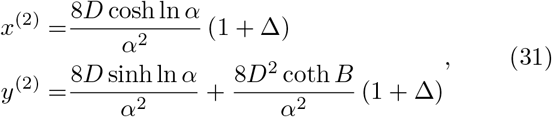

where

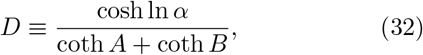

and *D* is invariant under the transformation 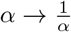. The second-order expression is then

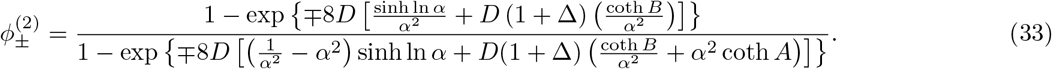

Notice again that the denominator is invariant under the transformation 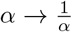. The second-order analytical form approximates the exact binodal to a high degree (figure 2B, C) even at the large-*N* regime.

## V. SELF-CONSISTENT EXPANSION

Although the self-consistent solutions are exact near critical points and convergent at large *χ*, the convergence is slow in the transition region. Here we show that we can improve the maps by performing a first order expansion of the self-consistent operator. Starting from a general self-consistent equation *η* = *𝒜* (*η*) and an initial guess *η*^(0)^ we want to find a step *δη* such that the next guess *η*^(1)^ ≡ *η*^(0)^ + *δη* solves the self-consistent equation to first order. We thus write *η*^(0)^ + *δη* = *𝒜* [*η*^(0)^ + *δη*]. Expanding *𝒜*[*η*^(0)^ + *δη*] to first order and solving for *δη* we obtain

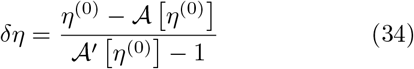

so the next best guess is

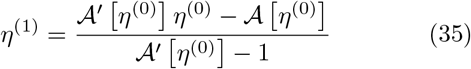

The above results can readily be applied to improve the maps *g*(*ϕ*) and ***G***(***ϕ***). In the *N* = 1 case we define the new map *h*(*ϕ*) as

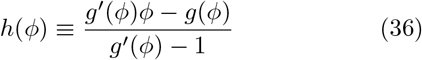

and it reduces to the original map when *g*′(*ϕ*) = 0. We thus define the new orbit 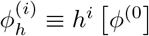. The convergent property can be studied by expanding the above with 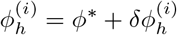, we arrive at

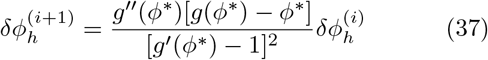

and near the fixed point the numerator approaches 0, so the convergence is rapid (figure 3A).

**FIG. 3.**
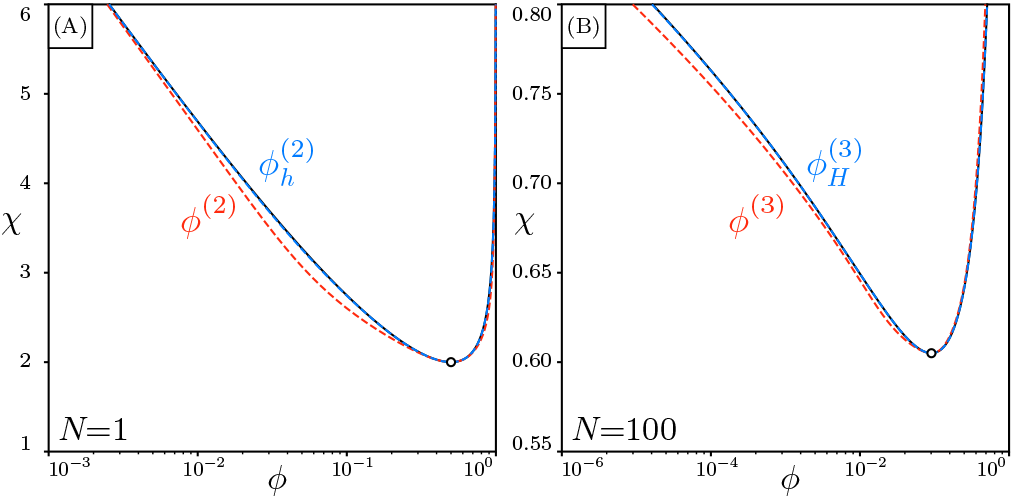
Improved self-consistent solutions from self-consistent expansion (blue dashed lines) converge to the true solution more quickly than the original ones (orange dashed lines), in both the (A) symmetric *N* = 1 case and (B) the general *N* case. The numerical binodal is black solid line and they virtually overlap with the blue dashed lines.

In the case of general *N* we similarly obtain

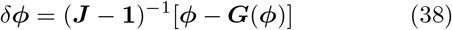

with **1** a 2 by 2 identity matrix. The improved operator is

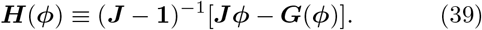

Good agreement with umerical results is achieved for the new orbit 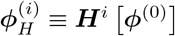 within three iterations for a large *N* = 100 (figure 3B). The self-consistent solution is thus able to cover a wide range of *N* with very few iterations, and can be used to perform numerical fitting of experimental binodal data.

## VI. DIRECT DATA-FITTING

The self-consistent solution allows efficient computation of binodal concentrations and we use it to fit experimental LLPS data and extract the interaction parameters. In the following fitting we use equation (33) to compute the binodal concentrations. Binodal concentrations for the prion-like low-complexity domain from isoform A of human hnRNPA1 (A1-LCD) were measured in [27] (137 amino acid residues) and three series of A1-LCD variants are fitted here. The first series involves aromatic residues Tyrosine (Y) and Phenylalanine (F); the second series involves non-equivalent polar spacers Glycine (G) and Serine (S); and the third series involves ionic residues Aspartic acid (D), Arginine (R) and Lysine (K). The wild type (WT) A1-LCD binodal was also measured.

### Fitting results

The chain length *N* is set as a global fitting parameter. The interaction parameter *χ* has the form 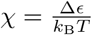 and we fit Δ*ϵ* for each variant. To convert concentrations to volume fractions we use a protein density of 1.35g/cm^3^ and average molecular weight of 13.1kDa to obtain the conversion ratio from concentration *c*(M) to volume fraction *ϕ* as 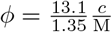. The fitting results give the effective chain length *N* = 158.6, larger than 137, the number of residues. This is as expected as *N* represents the number of lattice sites occupied by each polymer, and since the lattice site volume is determined by the underlying medium, i.e. water, the effective *N* will be larger than the number of residues owning to the larger size of amino acids compared to water molecules. The ratio 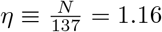 then represents the average number of lattice sites occupied by one residue. The fitted Δ*ϵ* values represent the site-to-site contact energy and a larger Δ*ϵ* indicates stronger attraction between proteins. To highlight the difference across variants we first calculate the protein-to-protein contact energy *ϵ* ≡ *−N* Δ*ϵ* and define the deviation from WT as Δ*E*_variant_ ≡ *E*_variant_ *− E*_WT_. Fitted curves are plotted in figure 4 and Δ*E* results are listed in table I. Each variant series then allows quantitative interpretation of impacts of different residues on LLPS propensity.

**FIG. 4.**
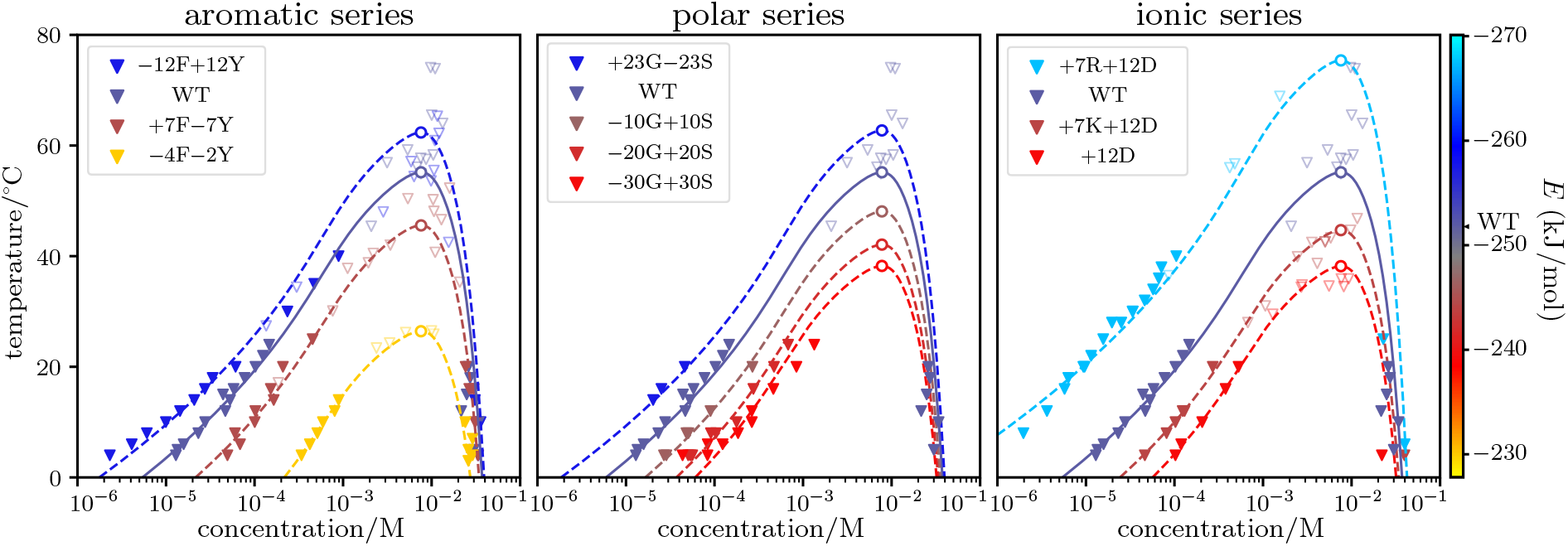
Flory-Huggins fit of binodal data from [27], fitted using equation (33). A constant *N* is maintained across all variants. Solid triangle markers are dilute and dense phase concentration measurements, and light hollow triangle markers are estimates of the critical point from cloud point measurements. Dashed lines are the best-fit curves and hollow circles are critical points. The WT binodal is the same in all three plots and is plotted in solid line. Colours of the plot represent the *E* ≡ *−N* Δ*ϵ* values of the variant. The *±nX* in variant names indicate *n* of *X* residues are added (+) or removed (−) from WT.

**TABLE I.** Fitting results for the A1-LCD data.

| aromatic series |  | polar series |  | ionic series |  |
| --- | --- | --- | --- | --- | --- |
| variant | $\Delta E$ (kJ/mol) | variant | $\Delta E$ (kJ/mol) | variant | $\Delta E$ (kJ/mol) |
| −12F+12Y | −5.5 | +23G−23S | −5.8 | +7R+12D | −15.5 |
| +7F−7Y | +7.4 | −10G+10S | +5.4 | +7K+12D | +8.0 |
| −4F−2Y | +22.0 | −20G+20S | +10.0 | +12D | +13.0 |
|  |  | −30G+30S | +13.0 |  |  |

### Tyrosine is a stronger sticker than Phenylalanine

In the aromatic series the effective interaction energy agrees with the observation that Tyr residues are stronger stickers than Phe residues [27], and from the fitted value we can further infer the individual contribution of each Tyr and Phe residue. Denote individual residue contributions as Δ*E*_Tyr_ and Δ*E*_Phe_, values from table I allow us to write down the following system of equations

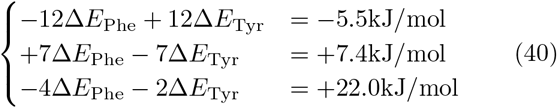

which can be solved to give Δ*E*_Tyr_ ≈ −3.2 kJ/mol and Δ*E*_Phe_ ≈ − 3.9 kJ/mol. Both Tyr and Phe are thus stickers with Tyr stronger than Phe.

### Serine destabilises condensates

Performing calculations similar to above we extract the difference Δ*E*_Gly_ − Δ*E*_Ser_ ≈ − 0.41 kJ/mol, indicating destabilising effect of Serine residues. This can be understood as the OH group in Ser forming favourable interactions with water, thus destabilising the condensate.

### Arginine promotes LLPS better than Lysine at constant charge

The ionic series data is harder to interpret since the overall protein charge can have a non-monotonic effect on LLPS propensity [27]. We can however still compare the +7R+12D and +7K+12D variants since they have the same overall charge. The energy difference between Arginine and Lysine is Δ*E*_Gly_ − Δ*E*_Ser_ ≈ − 3.4 kJ/mol, indicating stronger sticker behaviour for Arginine. This can arise due to the electron delocalisation in the guanidinium group and higher charge-charge contact efficiency with other charged residues.

## VII. CONCLUSION

We have developed a self-consistent solution for the binodal concentration of the 2-component Flory-Huggins phase-separating system. The proposed self-consistent operators shed light on the scaling behaviour of the dilute phase binodal, which is qualitatively different from the scaling of the spinodal and explains why LLPS of proteins occur over a concentration range spanning several orders of magnitude. Using the well-known Ginzburg-Landau binodal approximate solution as the initial guess, the self-consistent solution achieves numerical accuracy within 2 to 3 iterations and allows highly efficient fitting of experimental binodal data. Explicit analytical forms of the binodal concentrations are also proposed to approximate the binodal within one self-consistent iteration. Using the developed solution we fitted experimental data measured for variants of the A1-LCD protein and extracted effective interaction energies, which can be used to further decode the impact of individual amino acid residues on LLPS. Our analytical solution to the Flory-Huggins model thus allows systematic investigation of sequence grammar of LLPS-prone proteins and can lead to development of a wholistic framework for predicting LLPS propensity from sequence information.

## Code availability

A python library containing the self-consistent maps and auxiliary functions can be found at: https://github.com/KnowlesLab-Cambridge/FloryHuggins

## Acknowledgement

This study is supported by the Laboratory for Molecular Cell Biology, University College London (T.C.T.M.), the European Research Council under the European Union’s Seventh Framework Programme (FP7/2007-2013) through the ERC grant PhysProt (agreement no. 337969) (T.P.J.K.), the BBSRC (T.P.J.K.), the Newman Foundation (T.P.J.K.), and the Wellcome Trust Collaborative Award 203249/Z/16/Z (T.P.J.K.). The authors thank Alexander Röntgen for helpful discussions.

